# Mutation in SLC45A2 leads to loss of melanin in parrot feathers

**DOI:** 10.1101/2023.08.29.555428

**Authors:** Shatadru Ghosh Roy, Anna Bakhrat, Moty Abdu, Sandra Afonso, Paulo Pereira, Miguel Carneiro, Uri Abdu

**Affiliations:** Department of Life Sciences, Ben-Gurion University of the Negev, Beer Sheva 84105, Israel; STlab Hashita 240 Sede Tzvi, Israel; CIBIO, Centro de Investigação em Biodiversidade e Recursos Genéticos, InBIO, Universidade do Porto, Vairão, Portugal. BIOPOLIS Program in Genomics, Biodiversity and Land Planning, CIBIO, Vairão, Portugal

**Author notes:** **Corresponding author:** Uri Abdu, **Email:**.

**Keywords:** Feather, melanin, SLC45A2, Parrot

## Abstract

Bird plumage coloration is a complex and multi-factorial process that involves both genetic and environmental factors. Diverse pigment groups contribute to plumage variation in different birds. In parrots, the predominant green color results from the combination of two different primary colors-yellow and blue. Psittacofulvin, a pigment uniquely found in parrots, is responsible for the yellow coloration, while blue is suggested to be the result of light scattering by feather nanostructures and melanin granules. So far, genetic control of melanin-mediated blue coloration has been elusive. In this study, we demonstrated that feather from the *yellow* mutant rose-ringed parakeet displays loss of melanosome granules in spongy layer of feather barb. Using whole genome sequencing, we found that mutation in SLC45A2, an important solute carrier protein in melanin synthetic pathway, is responsible for the sex-linked *yellow* phenotype in rose-ringed parakeet. Intriguingly, one of the mutations, P53L found in yellow *P. krameri* is already reported as P58A/S in the human albinism database, known to be associated with human OCA4. We further showed that mutations in *SLC45A2* gene affect melanin production also in other members of Psittaculidae family such as alexandrine, and plum-headed parakeets. Additionally, we demonstrate that the mutations associated with the sex-linked *yellow* phenotype, localized within the transmembrane domains of the SLC45A2 protein, affect the protein localization pattern. This is the first evidence of plumage color variation involving *SLC45A2* in parrots and confirmation of associated mutations in the transmembrane domains of the protein that affect its localization.

**Article Summary:** The vibrant green tone of parrot plumage emerges from a blend of yellow and blue. While yellow comes from a unique pigment psittacofulvin, the genetics of blue structural color has puzzled scientists. Through novel genetic study, here we identified mutations in *SLC45A2* gene, causing loss of melanin in feathers that contributes to the blue structural color. This insight opens a new chapter in understanding avian coloration and the complex interplay of feather microstructure and genetics in nature’s palette.

## Introduction

Birds display one of the greatest ranges of coloration found in vertebrates. These striking colors are not just for our viewing pleasure. Every hue, stripe and spot serve a vital purpose. There are two main reasons behind it. First, birds use their colors to attract mates and intimidate competition. Secondly, they use them to provide protection from predators. Plumage coloration in birds can be created by several mechanisms. It can be the result of pigments, which absorb a certain wavelength of light, or the result of light scattering due to nanostructures present in the feathers (Prum et al. 1999). Parrots, known for their vibrant and diverse appearances, employ a combination of melanin, structural colors, and a unique pigment to achieve their striking plumage. While most birds utilize carotenoid pigments obtained from their diet for yellow and red coloration, parrots have a different approach (McGraw and Nogare 2004). Despite absorbing carotenoids from their food and circulating them in their bloodstream, parrots do not deposit these pigments directly into their feathers. Instead, they utilize a distinct group of yellow and red pigments called psittacofulvins (McGraw and Nogare 2004). However, the genetic mechanisms behind psittacofulvin production remain poorly understood. In 2017, an important breakthrough came when Cooke et al. (Cooke et al. 2017), studying captive budgerigars (Melopsittacus undulatus), identified a gene responsible for yellow psittacofulvin production. Through their research, Cooke and his colleagues demonstrated that a single amino acid substitution in an uncharacterized Polyketide synthase 1-like gene (MuPKS) disrupts the biochemical pathway for psittacofulvin synthesis. This genetic alteration is responsible for the “blue” trait observed in budgerigars. On the other hand, it has long been recognized that parrots exhibit reflective structural coloration in the short-wavelength range, specifically in the UV-blue spectrum (Prum et al. 1999; Hausmann et al. 2003). The structural coloration observed in parrots is heavily dependent on the microstructure or nanostructure of their feathers, including factors such as layering, ordering, particle sizes, and refractive indices (Prum et al. 1999). Similar to other organisms, the UV-blue plumage colors in parrots are generated by a specific microstructure within the feather tissue that leads to constructive interference of coherently scattered light waves (Prum et al. 1999). The spongy medullary layer of the feathers contains cells with quasi-ordered arrays of β-keratin rods and air vacuoles, with multiple such cells present in the ramus of each feather barb (Prum et al. 1999). The precise arrangement of these keratin rods at the channel type nanostructural level appears to influence the dominant wavelengths and saturation of the reflected light. Research on the production mechanisms of structural plumage coloration has been extensive over the past few decades, particularly through electron microscopy examinations (Prum 2022). These studies revealed that, the presence of feather nanostructure leading to coherent scattering seems to be widespread in non-cacatuid parrots, with blue or green coloration resulting from this coherent scattering observed in all parrots except cockatoos and eight species of Psittacidae or Loriidae (Eaton and Lanyon 2003). Cockatoos, in contrast, do not display UV-blue structural coloration based on coherent scattering, which explains the absence of green coloration in this group (Nemesio 2001). However, certain cockatoo species exhibit white plumage, generated by a feather structure that causes incoherent scattering of all visible wavelengths in the absence of pigments (Prum 2022). Notably, it has been shown that feathers from different budgerigar morphs, including grey and chromatic (purple to yellow) colors contained quasi-ordered air–keratin ‘spongy layer’ matrices, but these were highly reduced and irregular in yellow feathers. Similarly, yellow feather lacked a layer of melanin-containing melanosomes basal to the spongy layer. This data first suggested that the presence of melanosomes along with spongy layer might have role in production of blue color whereas their absence may allow solitary expression of yellow color (D’Alba, Kieffer, and Shawkey 2012). Melanin is presumably responsible for the grey-black plumage coloration displayed by many species, and electron microscopy has revealed the presence of melanin granules in pigmentary and structurally colored feathers. However, pigment in feathers exhibiting colors suspected of being produced by pheomelanin (i.e., dull red, yellow, greyish-brown and greenish brown) was tested using Raman spectroscopy in 26 species from the three main lineages of Psittaciformes. Non-sulphurated melanin form (eumelanin) in black, grey and brown plumage patches, and psittacofulvins in red, yellow and green patches were detected, but there was no evidence of pheomelanin (de Oliveira Neves, Galván, and van den Abeele 2020).

After decades of captive breeding, birds belonging to the order Psittaciformes, have surpassed their wild ancestors in terms of color diversity (van der Zwan and van der Sluis 2021). The range of color morphs exhibited by parrots now encompasses a huge number. The formation of these altered colors is believed to be influenced by a combination of nanostructures and pigments (Parker 2002). Given the abundance of color variations and the wealth of genetic information available from breeders, domestic parrots present an ideal model system for unraveling the underlying physical and genetic mechanisms behind avian color evolution. In this study, we employ microscopic as well as molecular tools to elucidate the physical mechanisms and genetic control responsible for color production in one of the distinct sex-linked morphs, known as *lutino* or *ino*, where melanin based grey or non-iridescent blue color is absent. We found mutation in *SLC45A2* is responsible for *yellow* phenotype in multiple members of Psittaculidae family. Our study also revealed that SLC45A2 fails to maintain its normal localization in the presence of mutations in its transmembrane domain.

## Materials and method

### Feather samples and light Microscopy

Contour and remex feathers of different color morphs of *Psittacula krameri* (Rose-ringed) were obtained from local breeders and pet shops. Feathers were washed in distilled water, dried overnight at 60°C and prepared for light microscopy. Images were taken using Leica M165 FC microscope.

### Feather histology

Parts of contour and remex feathers were washed in PBS, fixed in 4% paraformaldehyde overnight and washed in 50%-100% ethanol followed by xylene. After dehydration, the tissues were incubated in 65°C hot paraffin baths in vacuum oven for 1-2 hour each. Then, the samples were embedded in paraffin mold and sectioned at 8mm. Cross sections were observed and imaged under Nikon light microscope.

### Scanning electron microscopy

For scanning electron microscopy (SEM), parts of the feather were cut and stored in 1.5 mL microtubes. The samples were rinsed at least three times in distilled water. After that, samples were air dried, cut in smaller pieces using fine scissors and mounted on stubs with double-faced carbon cello-tape. Finally, sputtered with gold of 20nm depth before observation. The specimens were examined with a scanning electron microscope (Verios XHR 460 L).

### Sequence alignment

Homologs of *SLC45A2* were identified using the *P. krameri* GenBank assembly GCA_002870145.1. Identity was calculated via the Basic Local Alignment Search Tool (BLAST) (https://blast.ncbi.nlm.nih.gov/Blast.cgi). For pairwise sequence alignment EMBOSS Needle (https://www.ebi.ac.uk/Tools/psa/embossneedle/) was used. To identify the conserved sequences, a domain search was done using Jalview version 2.11.2.0 (Tcoffee alignment).

### Blood sample collections

Whole blood from a clipped toe was collected and dried on an appropriate filter paper with respective details from healthy birds owned by breeders. The collection was done by Sde-tzvi lab (https://dnalab.co.il/) as part of their sex identification services. The number of specimens that were examined by sanger sequencing are as follows: two wild type green, two *albino* (white) and twelve sex-linked *yellow* mutants from species *P. krameri*; two wild type green and ten sex-linked *yellow* mutants from species *Psittacula eupatria* (Alexandrine) and two wild type green and five sex-linked *yellow* mutants from species *Psittacula cyanocephala* (Plum-headed). Mutants examined in this study are shown in Figure 1.

**Figure 1:**
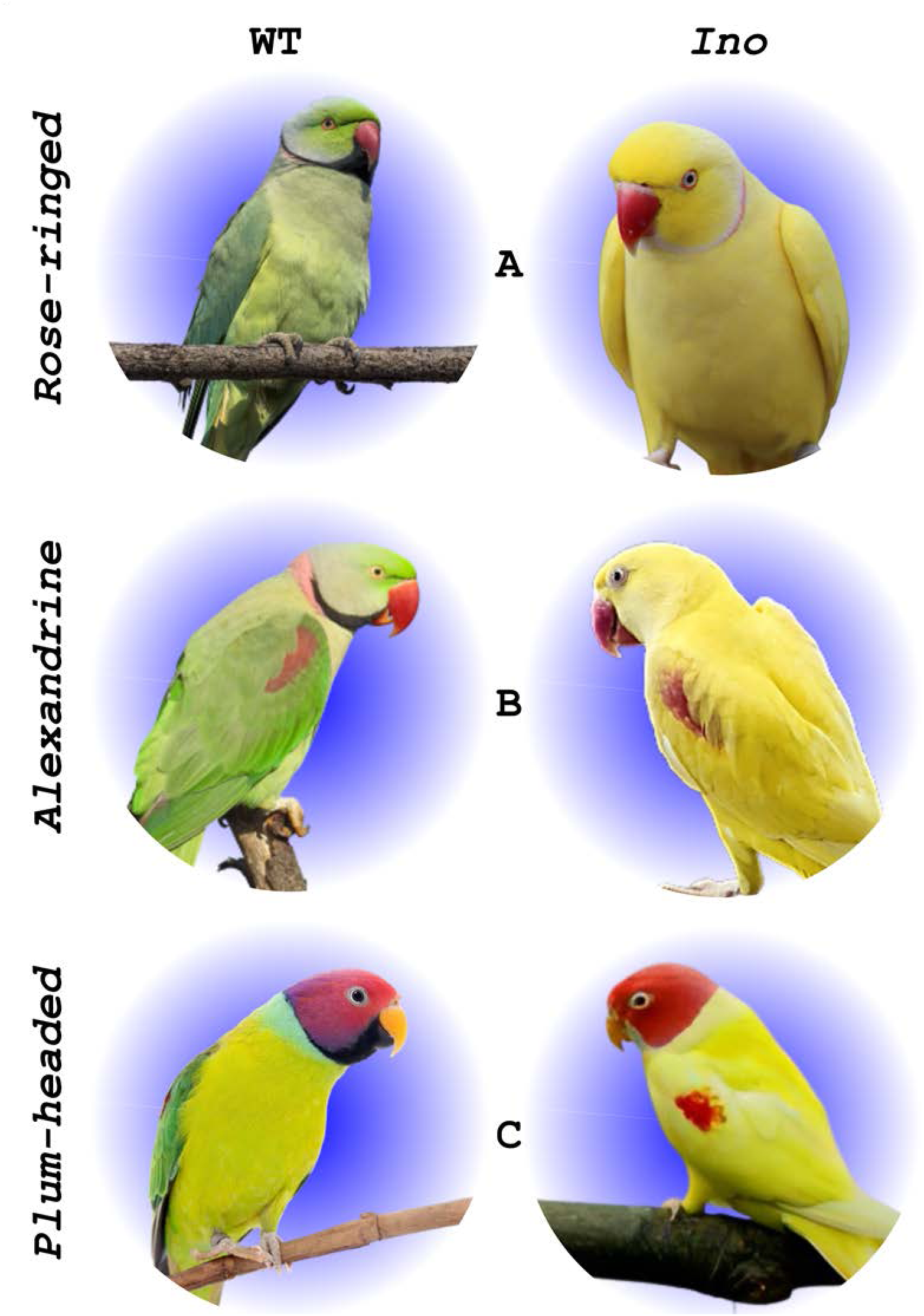
Species examined in this study from family Psittaculidae. A) Rose-ringed *(P. Krameri*); B) Alexandrine (*P. eupatria*); C) Plum-headed *(P. cyanocephala*). Wild type birds are at the left and *yellow* mutants are at right. (Pictures are taken either by Shatadru Ghosh Roy or available in public domain)

### DNA extraction from whole blood

Total DNA was extracted from whole blood using alkaline extraction method. Part of the paper with dried blood was soaked in 45ul 0.2M NaOH solution, followed by incubation at 75^0^C for 5 minutes. 1ul from this solution was used as template for PCR amplification mixture of 30ul.

### Whole-genome sequencing

DNA-seq libraries were prepared at the Crown Genomics institute of the Nancy and Stephen Grand Israel National Center for Personalized Medicine, Weizmann Institute of Science. 1ug of genomic DNA was mechanically sheared using a Covaris E220X ultra-sonicator to yield fragment size of ∼300 bp. Sheared DNA was run on TapeStation (Agilent) to ensure accurate shearing. Libraries were constructed as previously described (Blecher-Gonen et al. 2013) with the following modifications: end repair was done using 20U of T4 PNK and 6U of T4 Pol, and DNA cleanup was done with 0.75X Ampure beads. 10ul of 0.75uM adapters were used for adapter ligation. Eight cycles were used for the final library amplification. Libraries were quantified by Qubit (Thermo fisher scientific) and TapeStation (Agilent). Sequencing was done on a Nova-Seq 6000 instrument (Illumina) using a 300 cycles kit, allocating 150M reads per sample (paired end sequencing).

### Whole-genome sequencing data analysis

Sequence adaptors were trimmed with Cutadapt (Martin 2011) and mapped to the *Amazona guildingii* ASM1339961v1 assembly (PRJNA545868) using the BWA aligner (Li 2013) with default parameters. Technical duplicates were then excluded using the “UmiAwareMarkDuplicatesWithMateCigar” application in picard (https://broadinstitute.github.io/picard/; Broad Institute). Variant calling was carried out with FreeBayes (Garrison and Marth 2012). The coding sequence of genes of interest were converted from the reference to the variant form based on the called variants and the sequence was translated in order to evaluate the coding effect. Genes of interest were flagged where the character state of wild type green differed from that of *yellow* samples.

### Genotyping

Coding sequence of *SLC45A2* were amplified by PCR (2x PCR bios) from genomic DNA with intronic primers (Table S1) and Sanger sequenced with forward primer listed in tables. PCR product concentrations were determined by spectrophotometer at OD260 and ran on 1% agarose gel in TBE to validate its integrity. The genotypes for each bird were determined by manual inspection of the sequencing chromatograms.

### DNA constructs

For cloning HA tagged *PkSLC45A2* into pHAGE vector, firstly, total cDNA was prepared from total RNA, extracted from wild type green *P. krameri.* Coding sequence of *PkSLC45A2* gene was amplified by PCR and cloned into pRAT vector by Gibson assembly using SalI site. Subsequently, this coding sequence was sub-cloned into pPHAGE vector under CAG promoter with HA tag using KpnI site (used primers are listed in table S2). Nucleotide changes found in different species were introduced into pHAGE-HA-*PkSLC45A2* by site directed mutagenesis (used primers are listed in table S3).

### Cell culture and confocal microscopy

HeLa cells were cultured at 37°C and 10% CO_2_ in RPMI 1640 medium supplemented with 10% fetal bovine serum. Cells were transfected by PEI one day prior to imaging experiments with wild type HA-PkSLC45A2 and mutants. HA-*PkSLC45A2* transfected HeLa cells were fixed with 4% PFA for 15 minutes. The samples were then washed thrice in 0.3% PBST (Triton X-100) and incubated in 2% BSA in 0.3% PBST for an hour. After incubation with primary mouse anti-HA rat monoclonal antibody for 1hour (1:100; Sigma 11867423001) the cells were washed thrice with 0.3% PBST and incubated for an hour with secondary goat anti-rat Cy3 (1:200) antibodies (Jackson ImmunoResearch; 112-165-167). After washing thrice in 0.3% PBST, the samples were mounted in 50% glycerol. Images were collected with an Olympus FV1000 laser-scanning confocal microscope.

## Results

### Barbs of contour and remex feather exhibit different color patterns on the dorsal and ventral vanes

The typical pennaceous feather is comprised of a central shaft (rachis), with serial paired branches (barbs) forming a flattened or curved surface—the vane (Figure 2A). The barbs possess further branches called the barbules. A characteristic parrot feather can show the same or moderate to prominent color difference between its dorsal and ventral side, depending on its position. To examine the differential coloration of feathers from different parts, we carried out light microscopy for analyzing the vanes of two different types of feathers-remex and contour from both wild type and *yellow* mutant *P. krameri* (Figure 1). Remex is a flight feather found in the wings, while contour feathers form the bird’s outer body covering, giving shape and color to the bird (Prum and Brush 2002). Examination of a green remex from a wild-type *P. krameri*, showed green barbs on the dorsal side, with the barbules having yellow to black gradient towards the tip (Figure 2B). On the ventral side, barbs and barbules were uniformly colored as yellow (Figure 2C). Investigation of contour feather from wild-type *P. krameri* showed that the barbs were uniformly green on the dorsal side, while the barbules were yellow at the base and became black towards the tip (Figure 2D). However, at the ventral side, the barbs were green, and the barbules were entirely yellow (Figure 2E). Interestingly, light microscopic analysis of a contour and remex feather from the same species exhibiting sex-linked *yellow* phenotype displayed yellow barbs and barbules on both sides of the vane (Figure 2F-2I). Close examination of the barbules revealed that, in both types of feathers, dorsal barbules lost their black coloration at the tip, making it completely white in color (Figure 2F and 2H). Another noticeable difference was found at the ventral vane of remex feathers, where the yellow color of the barbs was markedly paler (almost white) than the ventral vane of the contour feather (Figure 2G). Overall observation implied that *yellow* trait abolishes blue structural color, leaving underlying yellow pigmentation on parts of dorsal and ventral vanes of feather, which otherwise appear green when blue structural colour is present.

**Figure 2:**
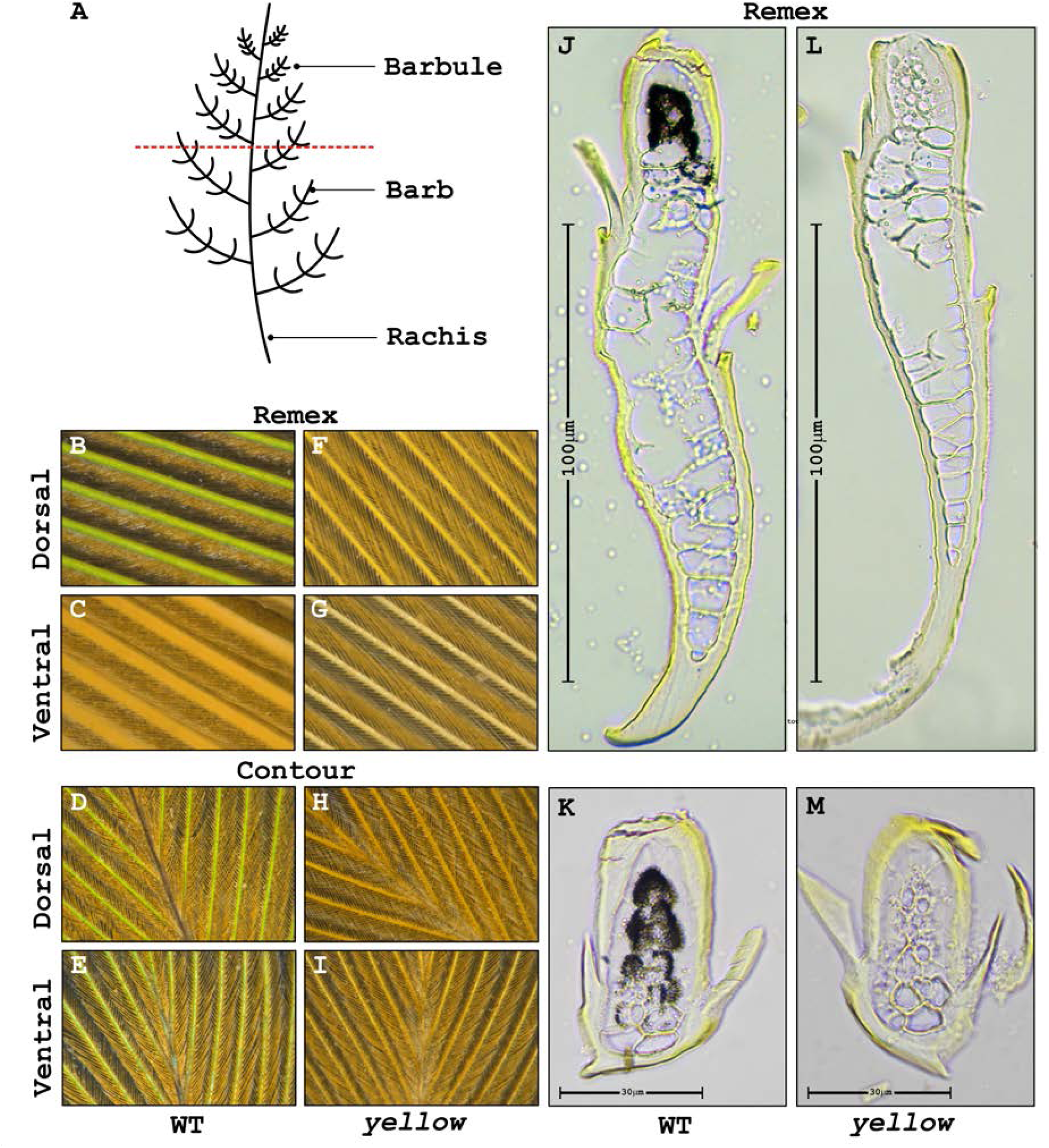
A) Diagrammatic representation of a typical pennaceous feather structure. B)-E) Panels show dorsal (B) and ventral (C) side of remex and dorsal (D) and ventral (E) side contour feathers from wild type *P. krameri.* F)-I) Panels show dorsal (F) and (G) ventral side of remex and dorsal (H) and ventral (I) side contour feathers from *yellow P. krameri.* J)-K) Light microscope images of cross-sections of the remex (J) and contour (K) feathers from wild type *P. krameri.* L)-M) Light microscope images of cross-sections of the remex (L) and contour (M) feathers from *yellow P. krameri.* Scale bars for light microscope images are 100 µm (remex) and 30µm (contour).

### *yellow* mutant displays loss of melanosome granules in spongy layer of feather barb

To characterize the microstructure responsible for the differentially colored feathers, we checked the cross-section of both remex and contour feather from green and yellow barbs under a light microscope. Cross-section of the green barb from a wildtype remex showed asymmetric organization from dorsal to ventral microstructure (Figure 2J). We found that the outer cortex is completely covered with yellow pigment and the dorsal side of the barb (upper part of the section in Figure 2J) carries melanin surrounding the central air vacuoles. Whereas the ventral side (lower part of the section in Figure 2J) of the barb is comprised of large air vacuoles across the medulla (Figure 2J). Cross-section of the green contour barbs from wild-type *P. krameri* showed two distinctive layers: 1) an outer cortex with yellow pigment around the entire periphery, and 2) a medullary region comprising of a layer of black melanin granules at the core, surrounding the central air vacuoles (Figure 2K). When we checked the cross-sections of contour and remex barbs from the *P. krameri* exhibiting the *yellow* phenotype, we found that barbs from *yellow* mutant feather share the same microstructural organization with the wild type green barb, comprising yellow cortex and spongy medullary region with central vacuoles (Figure 2L, 2M). But here, the black melanin layer at the core medulla is completely absent in both types of feathers (Figure 2L, 2M).

To confirm the arrangement of melanin granule and the structure of medullary layer, next we performed scanning electron microscopy (SEM) of cross-sections of contour and remex feather barbs from wild type and *yellow* mutants of *P. krameri*. In accordance with the light microscopy data, we observed in both wild-type remex and contour feathers, an outer cortex along with a medullary region comprises of large nuclear vacuoles. Nevertheless, SEM images confirmed the presence of spongy structure within medullary region together with several ovoid melanin granules around (Figure 3). A slight difference in length of the melanin granules was observed between remex and contour feathers. Where melanin granules of contour feathers are usually 1um in length, in remex feather the melanin granules were shown to be little longer (around 1.4um) in length (Figure 3B and 3F). The remex barb also showed polarized medullary spongy structure only at the upper part of the section corresponding the green dorsal side of the vane (Figure 3A). Together, SEM and light microscopy observations indicated that the green color of the vane is a combination of yellow pigment of the cortex with the blue from melanin granules through constructive interference of spongy medullary layer of the barbs. However, yellow vane is the result of the yellow pigment of the cortex alone, in absence of spongy structure or melanin granules embedded on it (Figure 3A, 3C). Therefore, no melanin granules were observed in spongy medullary region of yellow remex or contour feathers (Figure 3D, 3H). These results confirm that *the yellow* phenotype is characterized by a loss of melanin.

**Figure 3:**
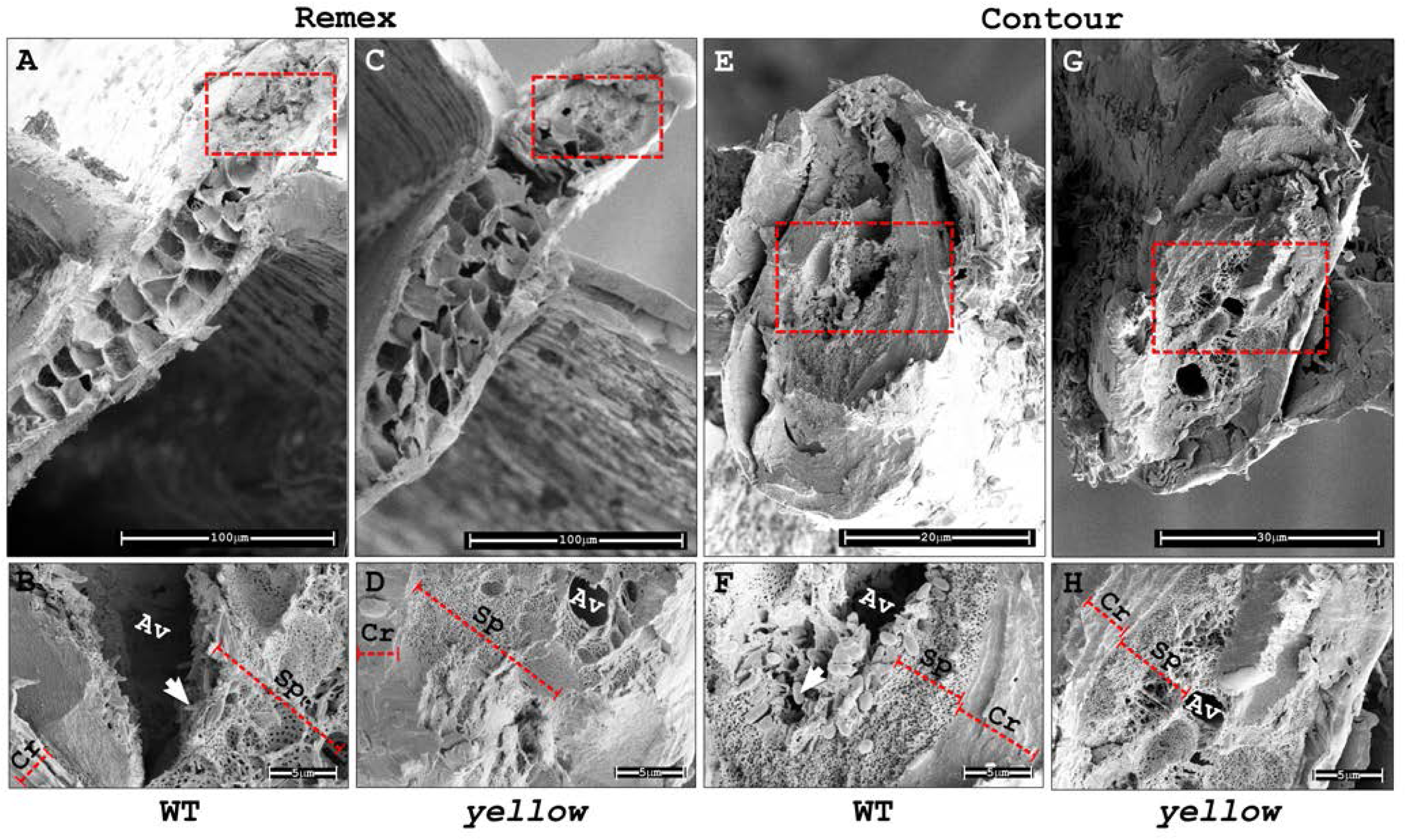
Scanning electron micrographs of remex feathers. A) Sectioned green barb of a remex feather from wild type *P. krameri.* B) Close-up view of a sectioned green remex barb, showing the outer cortex (Cr) with the layer of the spongy cells (Sp), with embedded thin elongated melanin granules (arrow). C) Sectioned yellow barb of a remex feather from *yellow P. krameri.* D) Close-up view of a sectioned yellow remex barb, showing the outer cortex (Cr) with the layer of the spongy cells (Sp), without melanin granules. E) Sectioned green barb of a contour feather from wild type *P. krameri.* F) Close-up view of a sectioned contour barb, showing the outer cortex (Cr) with the layer of the spongy cells (Sp), with ovoid melanin granules (arrow) embedded. G) Sectioned yellow barb of a contour feather from *yellow P. krameri.* H) Close-up view of a sectioned contour barb, showing the outer cortex (Cr) with the layer of the spongy cells (Sp), without melanin granules. Scale bars are as shown in the images.

### Identification of the *yellow* gene by screening single-nucleotide polymorphisms

The genetic control of avian melanogenesis is exerted by genes coding for specific enzymes involved in melanin synthesis and other important regulatory and structural proteins also needed for the process. In birds, more than 50 genes have been identified (Galván and Solano 2016). Most of these genes code for melanogenic hormones, receptors, factors for melanoblasts migration and melanocyte differentiation, structural proteins, and membrane transporters. Pedigree analysis shows that sex-linked *yellow* locus must reside on the Z chromosome. Therefore, we hypothesized that defect in melanin synthetic gene(s) located on Z chromosome are likely to be responsible for the *yellow* phenotype. Budgerigar genome data revealed that there are three annotated candidate genes present on Z chromosome. The first gene is *AGRP* (Agouti related neuropeptide), known to be an antagonist of the melanocortin-3 and melanocortin-4 receptor, which reverts melanogenesis towards the basal state of light pheomelanin synthesis from dark eumelanin synthesis (Tao 2010). The second is *TYRP1* (Tyrosinase related protein 1), a gene encoding a melanosomal enzyme from tyrosinase family, which functions in eumelanin production (Kobayashi et al. 1999), and the third one is a transporter protein that mediates melanin synthesis by transporting ions, named *SLC45A2* (Solute carrier protein 45A2) (Le et al. 2020). To identify candidate mutations responsible for the *yellow* phenotype in parrots, we sequenced the whole genomes of one green and two *yellow* mutants of *P. krameri*. The median coverage ranged between 30-60X. The coding sequence of the candidate genes described above were screened for protein-coding changes. Genes of interest were flagged where the character state of green differed from that of the *yellow* samples. The full list of flagged genes can be seen in file S1. After filtering out low-quality single-nucleotide polymorphisms (SNPs), significantly associated SNPs were found in a region predicted to be *SLC45A2* homolog of *yellow P. krameri* (hence named as *PkSLC45A2*). Two non-synonymous polymorphisms were identified at coding sequence of *PkSLC45A2* gene. One at the first predicted exon (c.188C>T) changing the 53^rd^ amino acid from proline to leucine and the next one at predicted exon six (c.1225G>A) changing the 399^th^ amino acid from glycine to arginine. No protein-coding changes were found in *AGRP* or *TYRP1*.

### Coding SNP in conserved PkSLC45A2 residue completely segregates with loss of pigmentation

To validate the mutations found in PkSLC452, we amplified the coding region of *PkSLC45A2* gene by PCR from genomic DNA of wild type green and mutants carrying the *yellow* phenotype. Sanger sequencing was done followed by pairwise alignment of the wild type and yellow mutant sequence data. The results were perfectly consistent with the expected segregation pattern. We found two expected non-synonymous missense variants c.188C>T; p.P53L and c.1225G>A; p.G399R in yellow mutants along with one new variant at exon 7 (c.1430G>T) changing 467^th^ amino acid from glycine to valine (Figure 4A, 4D). We used the same PCR primer of *PkSLC45A2* for amplification and sanger sequencing of *SLC45A2* homolog from two closely related parrot species, *P. eupatria* and *P. cyanocephala*, from the same Psittaculidae family (Figure 1). Comparison between *SLC45A2* sequence of green and *yellow* mutants revealed a missense mutation in *P. eupatria* at exon 2 (c.569T>C) changing 180^th^ amino acid from leucine to proline (Figure 4B, 4D), whereas a non-sense mutation at exon 1 c.103C>T in *P. cyanocephala*, altering 25^th^ amino acid arginine into a premature stop codon (Figure 4C, 4D).

**Figure 4:**
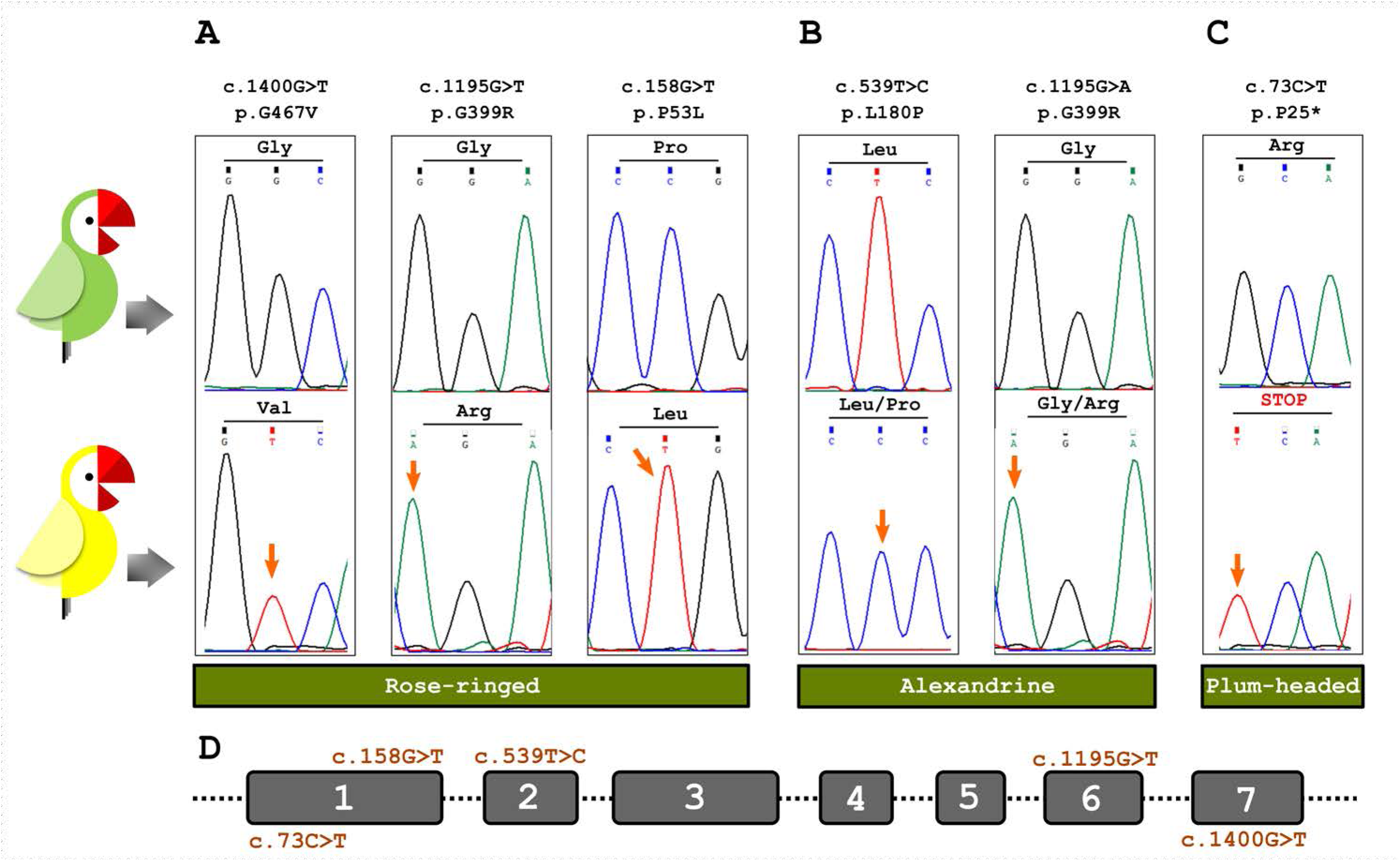
Non-synonymous protein-coding variations found in *SLC45A2* homologs. A) Missense mutations G467V, G399R and P53L found in rose-ringed (*P. krameri*). B) Missense mutations L180P and G399R found in alexandrine parakeet (*P. eupatria*). C) Nonsense mutation R25* found in Plum-headed parakeet (*P. cyanocephala*). D) Positions of nucleotide changes at different exons of SLC45A2 homologs.

A multiple sequence alignment showed that all the positions of the amino acid changes are universally conserved across fruit fly (*Drosophila melanogaster*), zebra fish (*Danio rerio*), chicken (*Gallus gallus*), gray short-tailed opossum (*Monodelphis domestica*), house mouse (*Mus musculus*) and human (*Homo sapiens*) (Figure 5A). Topology prediction data suggested that three of the non-synonymous polymorphisms i.e., P53L. L180P and G399R belong to important transmembrane domains 1, 5 and 9 respectively, whereas G467V was the only non-transmembrane variant, which was located at the penultimate cytosolic domain of the SLC45A2 protein.

**Figure 5:**
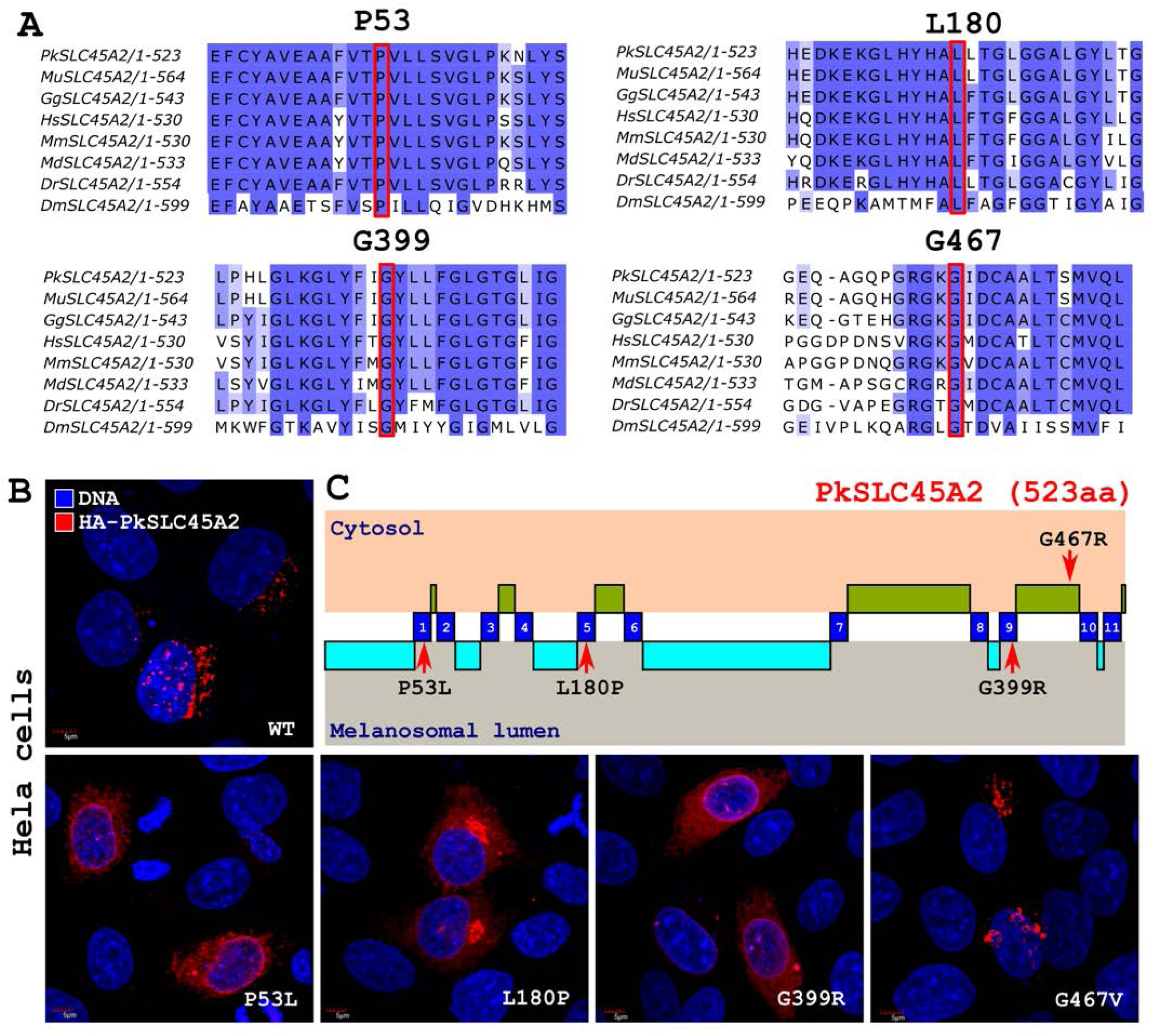
Mutation in the transmembrane domains of SLC45A2 affects the localization pattern of the proteins. A) Conservation status of changing amino acids found in *SLC45A2* homologs. B) Panel shows localization of wild type and mutant SLC45A2 upon ectopic expression in HeLa cells. C) Location of affected residues within important functional domains SLC45A2 protein. Transmembrane domains are depicted in blue, cytosolic part of the protein is in green and luminal part is in cyan.

### Mutations on the transmembrane domain affect localization of PkSLC45A2

To check any potential change in cellular localization pattern of wild-type and mutated *SLC45A2*, we expressed HA tagged PkSLC45A2 in HeLa cells under a constitutive promoter. In Hela cells, wild type HA-PkSLC45A2 was detected in distinct punctate microdomains associated with the nucleus. Next, we introduced the mutations (P53L, L180P, G399R and G467V) found in different parrot species into HA-*PkSLC45A2* plasmid by site-directed mutagenesis. All the mutations exhibited an altered pattern of localization except G467V (Figure 5B). HA-PkSLC45A2 with P53L, L180P and G399R mutations were detected to be dispersed throughout the cell when ectopically expressed. Interestingly, HA-PkSLC45A2 with G467V displayed the same punctate localization as wild type. Noticeably, three mutations with altered localization are located at different trans-membrane domains of the membrane associated transmembrane protein whereas, G467V mutation, showing same punctate localization as the wild type, resides at the penultimate cytosolic domain (Figure 5C). These findings suggest that transmembrane domains of SLC45A2 have a critical role in assuring its localization and maintaining its transporter ability.

## Discussion

### Barb microstructure plays crucial role in determining feather color

One key element of the green displayed in parrot feathers is blue structural color. Along with yellow psittacofulvin, selective absorption of blue wavelengths is necessary for the perception of green. Being unique in parrots, psittacofulvin has always been appealing to scientific attention. However, little is known about the exact mechanism behind non-iridescent blue structural color. The non-iridescent colors from the feathers of many birds are produced by quasi-random arrays of air vacuoles in the medullary keratin. Because they lack iridescence, these colors were hypothesized to be produced by differential scattering of light wavelengths by individual scatterers. Specifically, the colors were dominated solely by the scattering properties of individual air vacuoles instead of constructive interference of light (Tinbergen, Wilts, and Stavenga 2013). In early 70s, quasi-periodic photonic nanostructures were recognized in spongy barb medullary cells of parrot feather (Dyck 1971). Microscopic data revealed that blue structural color is generated by the spongy layer, which consists of nanoscale air cavities and β-keratin between the cortex and the medulla. Further research suggested that the presence or absence of melanin might also have a significant impact on blue color production (D’Alba, Kieffer, and Shawkey 2012). Initially, it was proposed that the lack of melanin in blue barbs results in a light blue color. However, studies have shown that melanin actually helps absorb scattered white light, preventing the washout of the coherently scattered blue color (Price-Waldman and Stoddard 2021). Feather barbs, except for yellow and grey ones, have a basal melanin layer underlying the spongy layer, indicating its critical role in non-iridescent structural color production. In addition, variations in the regularity of spongy medullary nanostructures contribute to different shades of blue, with higher regularity resulting in more saturated colors (D’Alba, Kieffer, and Shawkey 2012). In our study, we aimed to clarify the role of melanin distribution in the development of sex-linked *yellow* phenotype where structural blue color is completely missing. Using optical and scanning electron microscopy, we found that melanin granules in the medulla of green parrot feathers are oval and arranged in a palisade-like structure, embedded in the spongy medullary region. This structure was assumed to enhance the blue color reflection based on a simulation using Bragg’s formula (Okazaki 2020). Interestingly, the structure, size and the arrangement of melanin granule varied based on the feather type. We observed that the melanin granules of remex feather are thinner and longer than the same contour feather, which could be responsible for slight change in shades of green between two feathers. Additionally, we observed that barb microstructure devoid of spongy structure also failed to construct green color. In this study, we demonstrated that in *yellow* phenotype the feathers barb showed complete loss of melanin, suggesting defect in melanin synthetic pathway. In summary, our results implies that a characteristic parrot feather barb develops through keratin polymerization, melanosome synthesis and inclusion along with cortical psittacofulvin pigmentation. Compellingly, we found that even in the same bird, different feather types, based on its position varies through the microstructure of feather barbs, including the presence of psittacofulvin pigments, melanin granules, spongy medullary regions, and air vacuoles. Our study demonstrate that these components interact in complex ways to produce different colors in parrot plumage. The interplay between these components and their interactions with light influence the expression of different colors, offering insights into the mechanisms of color production in birds, in general.

### Defect in SLC45A2 function leads to loss of structural blue color

Sex-linked *yellow* phenotype are common in domesticated parrots. In our study, *yellow* mutants with absence of melanin were subjected to whole genome sequencing. The only gene found to have non-synonymous polymorphisms was *SLC45A2*. It produces a transporter protein found mostly in pigment cells. Changes in this gene can lead to a condition called oculocutaneous albinism type 4 (OCA4), which causes very little pigmentation (Le et al. 2020). These changes also contribute to differences in skin tone and skin aging in various human populations. In animals, equivalent gene mutations result in lighter pigmentation in species like gorillas (Prado-Martinez et al. 2013), dogs (Caduff et al. 2017), tigers (Xu et al. 2017), horses (Sevane, Sanz, and Dunner 2019), mice (Le et al. 2020), shrews (Tsuboi et al. 2009), chickens (Gunnarsson et al. 2007), quails (Gunnarsson et al. 2007), pigeons,(Domyan et al. 2014) frogs (Fu et al. 2022), and fish (Segev-Hadar et al. 2021). OCA4 patients have low pigmentation levels and show similarities to OCA2 patients who lack a melanosomal chloride channel (Bellono et al. 2014). This suggests that *SLC45A2* plays a significant role in melanogenesis (Fernández et al. 2021). The gene is primarily expressed in pigment cells and a few other cell types (Bin et al. 2015), as primary melanocytes from mice with the inactivating *underwhite* (uw) mutation of *SLC45A2* are hypopigmented, implying a melanocyte-depending defect (Newton et al. 2001). Although the exact function of the *SLC45A2* protein in melanocytes is not known, it bears some resemblance to sucrose transporters in plants and Drosophila (Newton et al. 2001; Meyer, Vitavska, and Wieczorek 2011), which are involved in transporting sugars. Pairwise alignment showed 62.43% of sequence identity of PkSLC45A2 with human SLC45A2 suggesting strong functional resemblance. Intriguingly, P53L mutation found in yellow *P. krameri* is already reported as P58A/S in the human albinism database (https://www.ifpcs.org/albinism/oca4mut.html), known to be associated with human OCA4.

When expressed in yeast, mouse *SLC45A2* functions at the plasma membrane as an acid-dependent importer of sugars (sucrose, glucose, or fructose) into the cytosol (Bartölke et al. 2014), suggesting that if SLC45A2 localized to acidic organelles it might facilitate export of a sugar and protons from the lumen to the cytosol. The key enzyme in melanogenesis, Tyrosinase, is inactive at pH < 6 (Halaban et al. 2002). So, neutralization of acidic early-stage melanosomes is a critical process for melanogenesis (Bellono et al. 2014). Consistent with a function in proton export and neutralization of acidic organelles, pigmentation of a zebrafish SLC45A2 mutant was rescued upon inhibition of endo-lysosomal and melanosomal acidification (Dooley et al. 2013). Moreover, knockdown of SLC45A2 in a pigmented melanoma cell line resulted in increased acidification of early-stage melanosomes (Bin et al. 2015). Interestingly, when *SLC45A2* is expressed in non-melanocytic cells, it localizes to late endosomes and lysosomes, similar to its distribution in some melanosomes in melanocytes. This indicates that SLC45A2 can also be found on lysosomes when expressed in other cell types and retains its ability to accumulate in specific membrane regions. In our study, we found that *SLC45A2* encoded membrane associated transporter protein with mutations at trans-membrane domain failed to maintain its original localization leading defective production of melanin granules. Overall, our data suggests that parrots carrying *yellow* phenotype lose their blue structural color due to defective expression of SLC45A2 in the melanocytes, that results in absence of melanin granules in the spongy medullary structure of the feather barb.

## Data availability

Plasmids are available upon request. DNA-seq data are deposited at GeneBank repository with accession numbers OR139606 (*P. krameri*), OR139607 (*P. eupatria*) and OR139608 (*P. cyanocephala*). Supplemental material is available at GENETICS online.

## Acknowledgments

We are grateful to Mr. Moty Abdu and Sde-tzvi lab (https://dnalab.co.il/) for providing the blood and feather samples. We would also like to thank you the Crown Genomics institute and the Mantoux Bioinformatics institute of the Nancy and Stephen Grand Israel National Center for Personalized Medicine, Weizmann Institute of Science, Israel. PP, SA and MC are supported by the European Research Council (ERC; to M.C.) under the European Union’s Horizon 2020 research and innovation programme (grant agreement No. 101000504).

